# Heart-brain interactions in the MR environment: characterization of the ballistocardiogram in EEG signals collected during simultaneous fMRI

**DOI:** 10.1101/185181

**Authors:** Marco Marino, Quanying Liu, Mariangela Del Castello, Cristiana Corsi, Nicole Wenderoth, Dante Mantini

## Abstract

The ballistocardiographic (BCG) artifact is linked to cardiac activity and occurs in electroencephalographic (EEG) recordings acquired inside the magnetic resonance (MR) environment. Its variability in terms of amplitude, waveform shape and spatial distribution over subject’s scalp makes its attenuation a challenging task. In this study, we aimed to provide a detailed characterization of the BCG properties, including its temporal dependency on cardiac events and its spatio-temporal dynamics. To this end, we used high-density EEG data acquired during simultaneous functional MR imaging in six healthy volunteers. First, we investigated the relationship between cardiac activity and BCG occurrences in the EEG recordings. We observed large variability in the delay between ECG and subsequent BCG events (ECG-BCG delay) across subjects and non-negligible epoch-by-epoch variations at the single subject level. Also, we found positive correlations between heart rate variability and ECG-BCG delay. The inspection of spatial-temporal variations revealed a prominent non-stationarity of the BCG signal. We identified five main BCG waves, which were common across subjects. Principal component analysis revealed two spatially distinct patterns to explain most of the variance (85% in total). These components are possibly related to head rotation and pulse-driven scalp expansion, respectively. Our results may inspire the development of novel, more effective methods for the removal of the BCG, capable of isolating and attenuating artifact occurrences while preserving true neuronal activity.

## 1. Introduction

The simultaneous acquisition of electroencephalography (EEG) and functional magnetic resonance imaging (fMRI) data enables the investigation of human brain function with high spatio-temporal resolution (Mantini et al., 2010). Simultaneous EEG-fMRI is nowadays widely used, particularly for the non-invasive identification of epileptic foci (Grouiller et al., 2016) and for the mapping of neuronal oscillations during sleep (McAvoy et al., 2017), rest (Mantini et al., 2007) or active task performance (Debener et al., 2005). However, the use of simultaneous EEG-fMRI remains technically challenging, especially due to the presence of fMRI-related artifacts in the EEG data (Neuner et al., 2014). These artifacts originate from the interactions between the subject, the EEG system and the magnetic MR environment. The two major artifacts that corrupt the EEG recordings are: the imaging artifact, which is caused by the time-varying magnetic field gradients applied during the fMRI acquisition (Debener et al., 2008); and the ballistocardiographic (BCG) artifact, which is associated with the cardiac activity of the subject (Allen et al., 1998).

Whereas the imaging artifact can be easily removed due to its periodicity and stability across time, the BCG artifact does not show such consistency. This poses important challenges for its attenuation from EEG data collected during fMRI scanning (Debener et al., 2007; Mullinger et al., 2013). Several studies were conducted to clarify the physical mechanisms at the basis of the BCG (Yan et al., 2010; Mullinger et al., 2013). They revealed that the BCG mainly arises from slight movements of the EEG electrodes and the wires in the static magnetic field, following each heartbeat, that produce an electromotive force that adds up to the EEG signals. In particular, voltage variations on the scalp are generated by the contribution of three main effects: 1) pulse-driven expansion of the scalp, i.e. the local motion associated with the pulsatile expansion of scalp vessels on adjacent electrodes (Bonmassar et al. 2002; Yan et al., 2010; Mullinger et al., 2013); 2) pulse-driven rotation of the head, i.e. and the motion derived from quick arrival and shunting of the blood into the head arteries (Yan et al., 2010); and 3) the pulsatile flow of blood that, as an electrically conducting fluid, in a magnetic environment, produces a separation of charges via Hall effect that induces the potential variations at the scalp surface (Müri et al., 1998). It is not clear, however, the relative contribution of these three components to the measured BCG, neither their spatial distribution across EEG sensors.

Several methods have been developed in the last years for BCG artifact removal, such as the adaptive average subtraction (AAS) (Allen et al., 1998), optimal basis set (OBS) (Niazy et al., 2005) and independent component analysis (ICA) (Mantini et al., 2007; Srivastavna et al., 2007). Each of these methods relies on specific hypotheses about BCG features, which need to be fulfilled for an effective artifact removal. Specifically, both AAS and OBS assume a fixed delay between the BCG and the associated cardiac events (Allen et al., 1998; Niazy et al., 2005). Also, ICA assumes the spatial stationarity of the sources (Krishnaswamy et al., 2016), such that the EEG recordings can be expressed by an instantaneous linear mixture of independent components. To the best of our knowledge, no study has so far been conducted to test whether such hypotheses for the BCG artifact effectively hold. On the other hand, considering that the BCG residuals left by any the aforementioned methods in the EEG data is not negligible (LeVan et al., 2014), we argue that the hypotheses may not be fully met.

In this study, we conducted a detailed characterization of the BCG ballistocardiogram in EEG signals collected during simultaneous fMRI. First of all, we investigated whether the delay between BCG and cardiac events is fixed, or is alternatively variable and dependent on the cardiac frequency. We then examined the spatio-temporal variations of this artifact both at the subject level and at the group level, to test whether the BCG sources can be considered spatially stationary. The findings of this study revealed novel insights into the BCG artifact properties. This was particularly important for identifying possible limitations of currently used BCG artifact removal methods and for inspiring the development of novel methods that overcome such limitations.

## 2. Materials and Methods

### 2.1.1. Data acquisition

Six right-handed healthy subjects (age 26.7±6.2 years, 4 males and 2 females) participated in the experiment. All participants reported normal or corrected-to-normal vision, and had no psychiatric or neurological history. Before undergoing the examination, they gave their written informed consent to the experimental procedures, which were approved by the local Institutional Ethics Committee of UZ Leuven.

Each subject underwent a 10-minutes resting-state session, during which EEG and fMRI data were concurrently recorded. fMRI imaging was performed for 10 minutes in a 3T Philips Achieva MR scanner (Philips Medical Systems, Best, the Netherlands) using a T_2_*-weighted SENSE sequence. The scanning parameters were TR = 2000 ms, TE = 30 ms, 36 slices, 80 × 80 matrix, voxel size 2.75 × 2.75 × 3.75 mm3, flip angle = 90 degrees. EEG data were acquired with a MR-compatible 256-channel HydroCel Geodesic Sensor Net (EGI, Eugene, Oregon, USA), that includes a large number of Ag/AgCl electrodes on lower temporal areas and cheeks. Electrodes impedance were kept below 50 kΩ across the full recording. An elastic bandage was also placed above the EEG net to maintain the contact of electrodes on the scalp. EEG signals were acquired at 1 kHz sampling frequency using the vertex (Cz) electrode as physical reference. Using the EEG amplifier, we also acquired the electrocardiogram (ECG) signal with two MR-compatible electrodes positioned on the chest, in correspondence to the apical part and to left side of the heart.

### 2.1.2. EEG data preprocessing

EEG data were processed by using built-in MATLAB (MathWorks, Natick, US) functions and the EEGLAB toolbox (https://sccn.ucsd.edu/eeQlab/) (Delorme and Makeig, 2004). First of all, the imaging artifact in the EEG and ECG data was attenuated by using the fMRI artifact template removal (FASTR) method implemented in EEGLAB (Niazy et al., 2005). The EEG signals were then digitally filtered in the frequency band [1-70 Hz], They were re-referenced in average reference (Liu et al., 2015), which allowed the reconstruction of the EEG signal at the Cz location. The re-referenced EEG data were finally processed for characterizing the BCG artifact and identifying its relation with the subject’s cardiac activity, as detected from the ECG signal.

### 2.1.3. Identification of ECG and BCG peaks

The ECG signal was band-pass filtered using a Finite Impulse Response (FIR) filter with a low and high cut-off frequencies of 5 and 20 Hz, respectively. The ECG peaks were detected identifying local maxima spaced in time by at least 600 ms. Following this automated detection, we manually corrected for false positives (less than 1.5% of the total) and negatives (less than 1% of the total). The detection of BCG peaks was performed using the EEG signal at the Cz location, which was standardized across subject during data acquisition. The EEG signal was filtered in the band [5-20 Hz], as for the ECG signal. BCG peaks were defined as the points with maximum cross-correlation with the average ECG signal. Since the BCG peaks were expected to follow the ECG peaks by around 200 ms, the selection of the points with maximum cross-correlation was restricted to the time window from 100 to 300 ms after the ECG peaks.

### 2.1.4. Characterization of BCG artifact

We examined the variability of the delay between ECG and BCG peaks, and their relation with heart rate variability (HRV). In particular, we evaluated the linear relationship between the duration of each cardiac period and the associated ECG-BCG delay, both at the single subject and group level.

We then examined the spatial stationarity of the BCG artifact, by calculating average EEG signals at the individual level, and using the ECG events as triggers. We identified and isolated in each subject corresponding BCG waves based on the root mean square (RMS) across channels. The scalp maps corresponding to the selected BCG waves were compared within and across subjects by using the Pearson correlation, to test for spatial stationarity of the BCG sources. In the case of stationarity, the correlations between different time instants for the same subject were expected to be larger than those between corresponding time instants for different subjects.

After identifying and isolating in each subject corresponding BCG waves, we also run PCA on concatenated data to examine the most prominent contributions to all BCG waves. Only principal components (PCs) with associated variance above 10 % were retained for analysis. For each selected PC, we plotted the associated topographic map to examine the spatial distribution across EEG channels.

## 3. Results

Imaging artifact removal was successfully performed in each EEG dataset (for an example, see Supplementary Figure 1). We identified ECG and BCG peaks, and extracted ECG-BCG delay and cardiac cycle duration accordingly (Figure 1). We could not obtain a reliable detection of the BCG events in one subject (S6), which was excluded from the current analysis. First, we analyzed HRV using the BCG peak information, and observed normal, physiological variability within and across subjects (Figure 2a,b and Table 1), with average values ranging between 843 and 1085 ms. Also the ECG-BCG delay showed a variability that was qualitatively similar to the HRV (Figure 2c,d). This figure was not only variable within each subject, but even more across subjects (Table 1). Specifically, average values ranged between 158 ms and 261 ms. The correlation between HRV and ECG-BCG peaks delay was positive but not always strong at the subject level (Table 1). Conversely, we obtained a robust positive correlation, which was equal to 0.47, when we used concatenated data across subjects (Figure 3).

**Figure 1.**
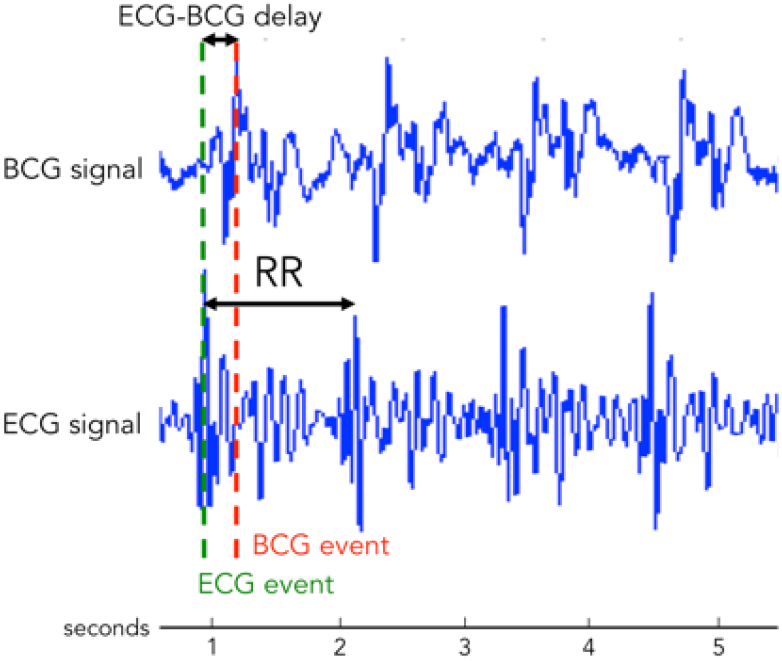
Detection of ECG and BCG events. Representative traces displaying the timing relationship (ECG-BCG delay) between the R-peak from the ECG signal (ECG event) and the main peak of the BCG occurrence (BCG event) from the EEG recordings acquired inside the MR scanner. The interval between two consecutive cardiac occurrences is defined as RR.

**Figure 2.**
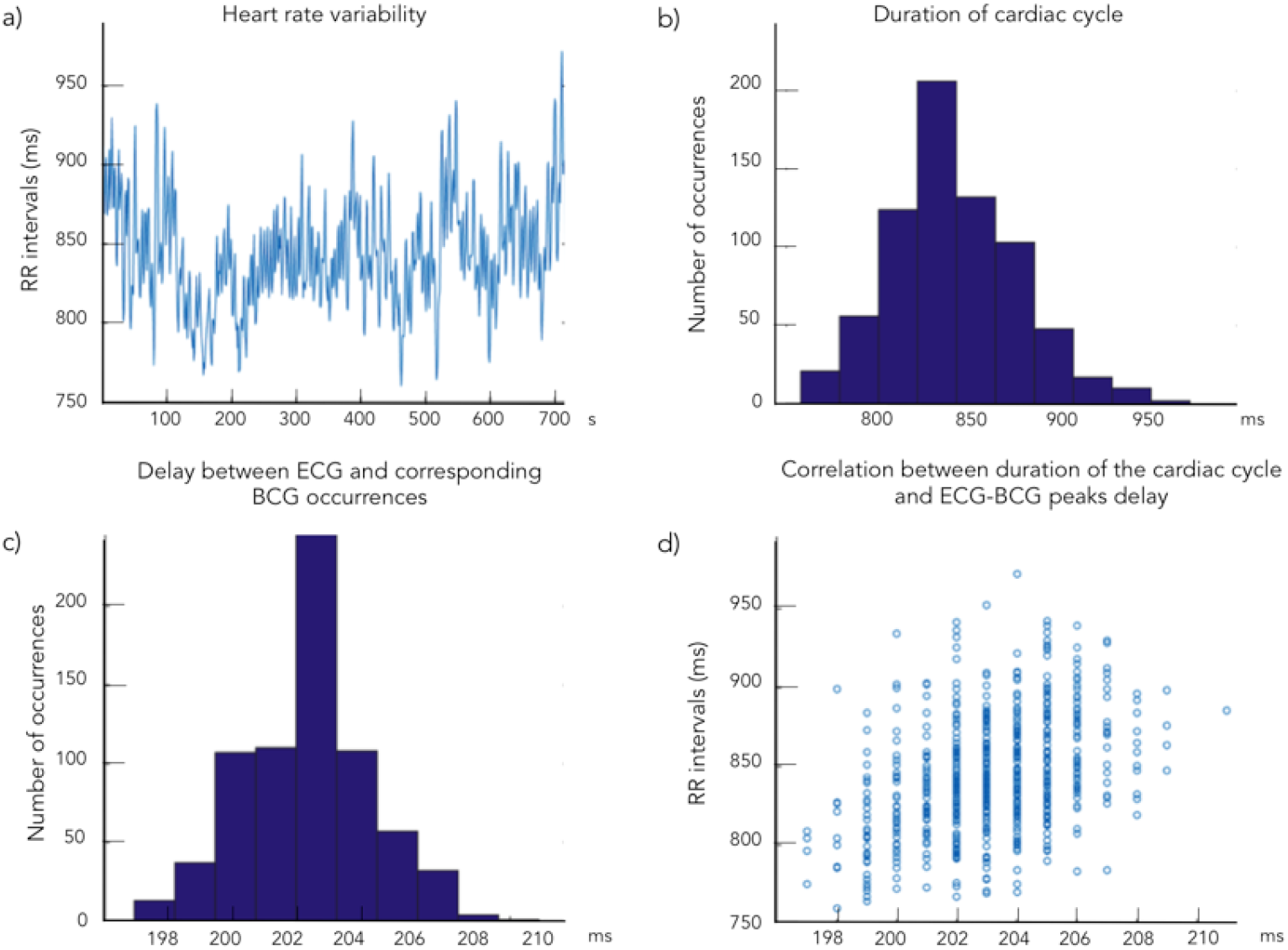
Heart Rate Variability (HRV) and ECG-BCG delay variability. a) The HRV is estimated by considering the sequence of RR intervals, i.e. time intervals between each ECG-peak and its consecutive, and b) the distribution of the RR intervals is calculated, c) The delays between each cardiac event and its corresponding BCG event are extracted and its distribution is calculated, d) The relationship between RR intervals and ECG-BCG delays are shown by means of a scatter plot.

**Figure 3.**
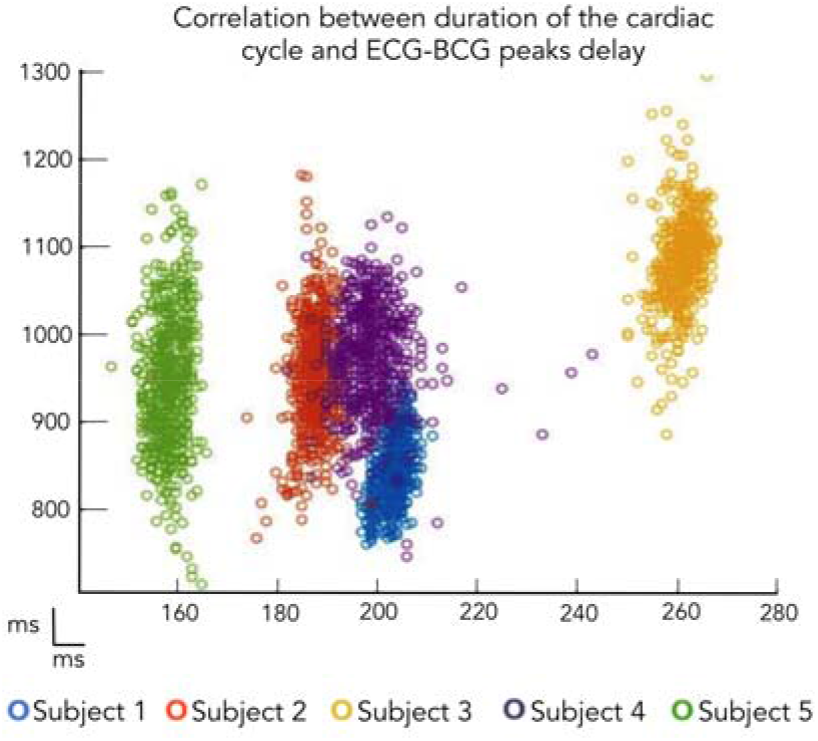
Relationship between RR intervals and ECG-BCG delay. The scatter plot shows a dependency between HRV and the ECG-BCG delay variability across subjects.

**Table 1.**
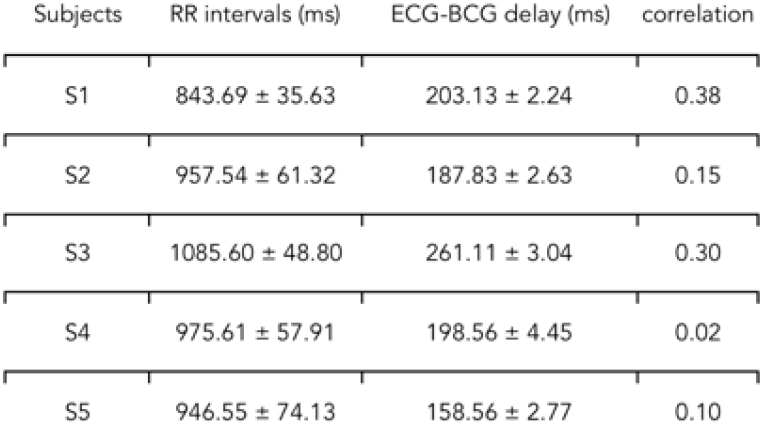
Descriptive statistics for RR intervals and ECG-BCG delays in individual subjects. The relationship between the two parameters was assessed by means of Pearson’s correlation.

| Subjects | RR intervals (ms) | ECG-BCG delay (ms) | correlation |
| --- | --- | --- | --- |
| S1 | 843.69 ± 35.63 | 203.13 ± 2.24 | 0.38 |
| S2 | 957.54 ± 61.32 | 187.83 ± 2.63 | 0.15 |
| S3 | 1085.60 ± 48.80 | 261.11 ± 3.04 | 0.30 |
| S4 | 975.61 ± 57.91 | 198.56 ± 4.45 | 0.02 |
| S5 | 946.55 ± 74.13 | 158.56 ± 2.77 | 0.10 |

After studying the variability in the ECG-BCG delay, we focused on the spatio-temporal characteristics of the BCG. We calculated averaged BCG signal using ECG peaks as events, also to allow comparisons with previous studies. An analysis of the RMS across channels confirmed the complex nature of the BCG artifact, revealing 5 major peaks that were present in all subjects at similar latencies (Figures 4 and Supplementary Figure 2). The non-stationarity of the BCG sources was indicated by the fact that the scalp maps for different time instants in the same subject had lower correlation (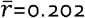; range [0.053-0.510]) than those of corresponding time instants for different subjects (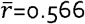; range [0.431-0.741]).

**Figure 4.**
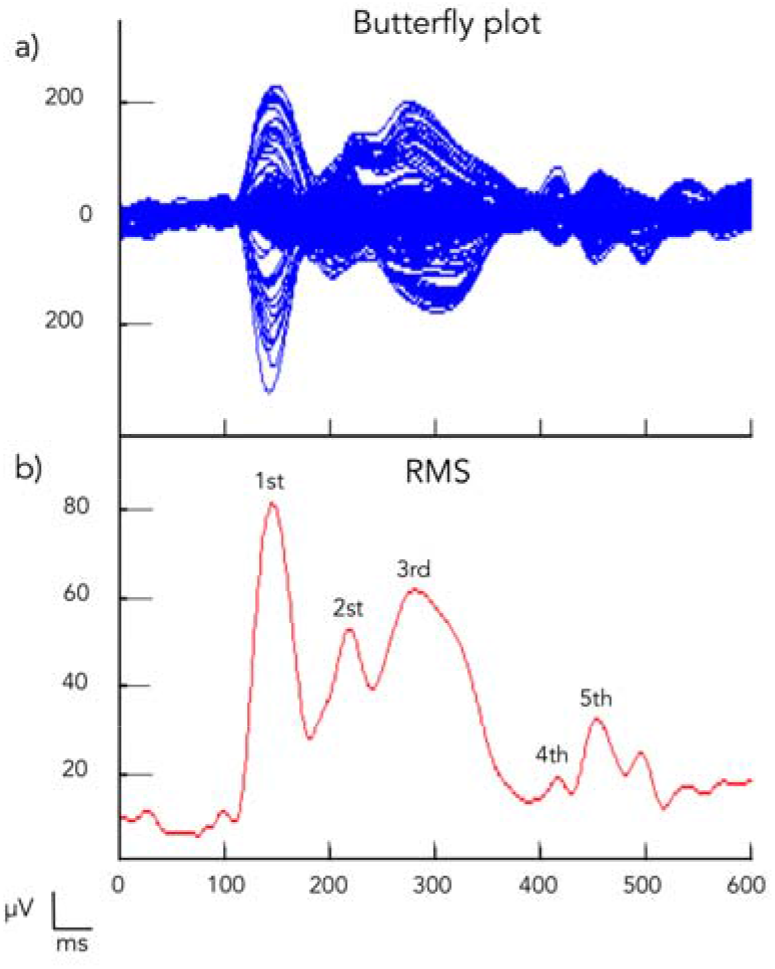
Selection of main BCG waves in individual datasets. a) EEG data were epoched by using the ECG events as triggers to produce a butterfly plot; b) RMS across channels was computed to identify five main peaks (marked in the figure).

By averaging the topographic maps of all the subjects, we retrieved characteristic scalp activity profiles at each major peak (Figure 5). In particular, we found that the BCG artifact was initially localized in the left side of the head at the first peak instant, and it progressively spreads out towards the right side. The application of the PCA enabled the identification of two main components, with associated variance equal to 76.69% and 12.71%, respectively (Figure 6). The first component was stronger on the left side of the scalp, whereas the second component was more diffused and bilateral. These two components had a spatial pattern very similar to the ones at the first and second peak instants, respectively, suggesting that the major effects of the BCG artifact occurrences were largely independent and concentrated at earliest latencies.

**Figure 5.**
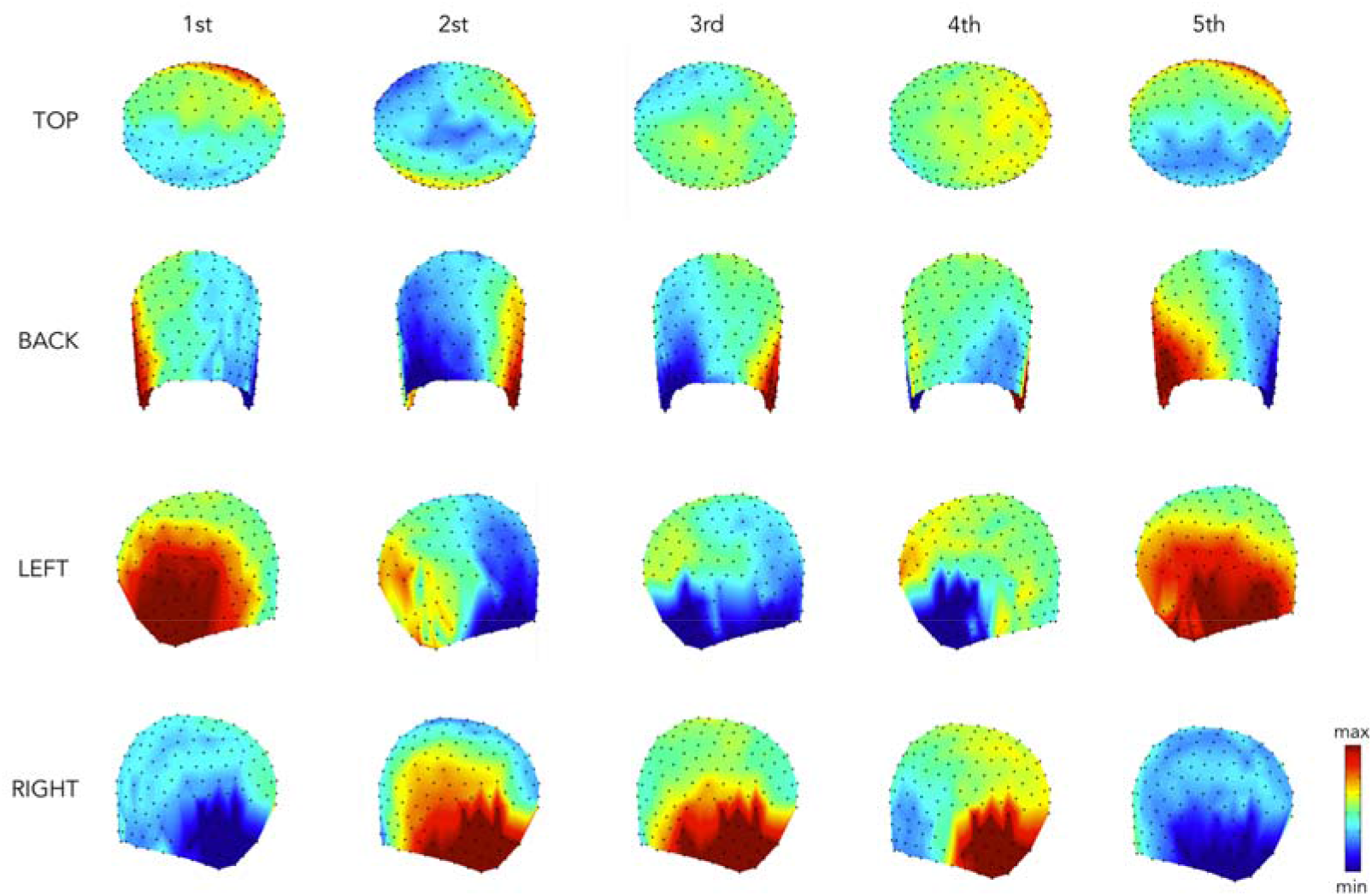
Group-level spatial maps for each of the five main BCG peaks. Topographic maps were extracted for each subject in correspondence to five different major peaks identified from the RMS plot. They were then normalized to z-scores and averaged across subjects.

**Figure 6.**
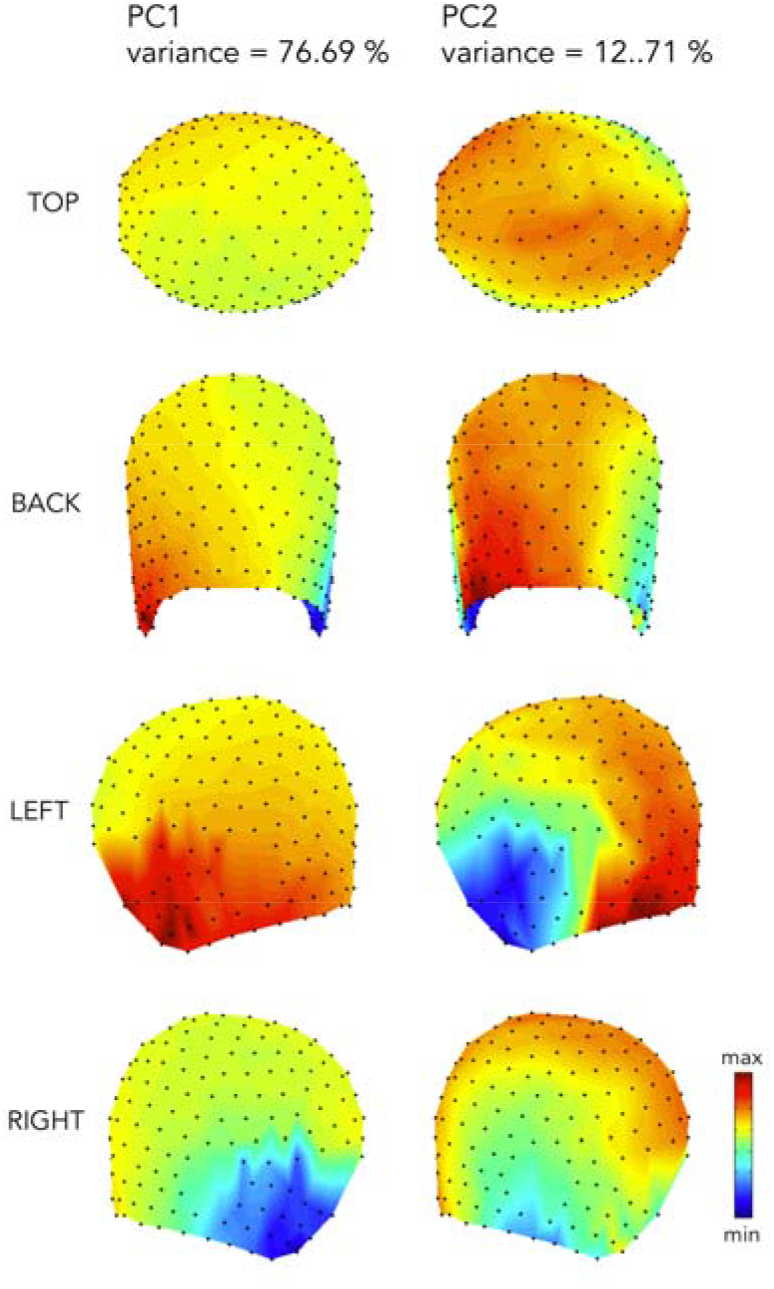
Spatial maps of the two main BCG components. The first (left) and second PCs (right) had 76.69% and 12.71 % explained variance, respectively. Their topographic maps are shown in z-scores for visualization purposes.

## 4. Discussion

In this study, we have investigated the BCG artifact and provided new insights concerning its relationship with cardiac activity and its spatio-temporal properties. Our analysis showed that BCG artifact occurrences follow each cardiac event with a variable delay (Figure 2 and Table 1). Furthermore, the ECG-BCG delay variability has a moderate positive correlation with HRV, i.e. when the cardiac frequency decreased the BCG artifact appeared on the EEG recordings with a smaller delay with respect to the cardiac event (Figure 2–3 and Table 1). Secondly, our analyses provided corroborating evidence for the non-stationarity of the BCG artifact (Figure 5). Specifically, we found two main components to be present in the BCG, with a specific spatial distribution and varying intensity across time (Figures 6).

### Variable delay between ECG and BCG peaks

Our study provided strong evidence for a variable delay between ECG and BCG peaks (Figure 2). This is in line with recent BCG artifact removal studies suggesting that accounting for a variable delay may enable a more effective artifact attenuation (Oh et al., 2014; Iannotti et al., 2015). It should be considered that our investigation on the delays between ECG and BCG peaks was conducted using the ECG signal and just one representative EEG signals, i.e. the one at Cz electrode. Indeed, some EEG channels do not show a clear BCG peak, which affects the reliability of peak detection. More importantly, we chose to use the EEG signal from the Cz electrode since the position of this channel corresponds to the head vertex, and is therefore standardized across subjects. This ensures that the differences observed across subjects are true differences, and do not depend on the different positioning of the EEG net. For the first time, our study showed a relationship between HRV and the ECG-BCG delay (Figures 2–3 and Table 1). This means that an increase in the duration of the cardiac cycle, i.e. decreased cardiac frequency, entails an increase in the ECG-BCG delay. A larger dataset would be desirable to provide stronger evidence for this heart-brain interaction effect. Also, further research into the underlying physiological mechanisms is warranted.

Interestingly, our analyses revealed that the latency variability of the BCG artifact occurrences was largely inconsistent across subjects (Oh et al., 2014; Iannotti et al., 2015). Accordingly, the use of a fixed delay between ECG and BCG peaks in BCG removal algorithm such as AAS (Allen et al., 1998) and OBS (Niazy et al., 2005), which is typically set to 210 ms, can be hardly justified. Furthermore, the intra-subject variability of the delay is a consistent feature of the BCG, and it should be taken into account for optimizing the removal of this artifact from EEG data. An accurate alignment of BCG epochs may indeed lead to improved performance when using AAS and OBS.

### Spatio-temporal analysis of the BCG

In order to study the spatio-temporal characteristics of the BCG, we calculated averaged EEG signals using ECG peaks as events. One may argue that this is not fully in line with the observation that the BCG peaks have variable delay with respect to ECG peaks. Our choice was primarily due to the fact that the same approach was used as in previous BCG studies (Debener et al. 2008; Mullinger et al., 2013). This would have enabled us to comparing our findings with those in the literature. Our analysis of the BCG showed a very heterogeneous spatial distribution overtime, suggesting that the artifact is non-stationary (Figures 4–5). This means that the use of BCG artifact removal methods that assume the stationarity of the sources, such as ICA (Comon, 1994), may not lead to optimal results.

Despite the complex spatio-temporal properties of the BCG, we were able to identify five main BCG waves, which were spatially consistent across subjects (Supplementary Figure 2). To this end, we used the RMS plot (Figure 4), which was already employed in previous studies (Debener et al., 2008). The scalp regions mostly affected by the artifactual contribution appeared to be the fronto-lateral and the occipital ones (Figure 5). A strong BCG in fronto-lateral part of the EEG montage is consistent with the presence of major blood vessels (Bonmassar et al., 2002; Masterton et al., 2007) and with the specific orientation of the electrodes with respect to the static magnetic field (Yan et al., 2010). A possible reason for a relatively strong BCG in the occipital part of the EEG montage is the adherence of the recording EEG electrodes on the subject’s head in the MR scanner. Previous studies suggested that the BCG may result from two distinct originating mechanisms, head rotation and pulse-driven expansion, respectively (Yan et al., 2010; Mullinger et al., 2013). In line with this literature, we experimentally found two main BCG components that are consistent across subjects (Figure 6). Based on the spatial distribution of these components, we argue that the first component (77% variance) may be associated with head rotation (Mullinger et al., 2013), whereas the spatial distribution of the second component (13% variance) may be compatible with pulse-driven scalp expansion. Future studies are warranted to verify the correctness of the suggested associations.

## 5. Conclusions

In this study we performed a detailed characterization of the BCG in EEG signals collected during simultaneous fMRI. This may contribute to a deeper understanding of the BCG spatial-temporal features. Specifically, we showed for the first time a relationship between HRV and variability in the delay between ECG and BCG events. We also identified two patterns that primarily contribute to the BCG, which have specific spatial and temporal features and can be therefore associated with different physiological sources. We hope that the findings presented in our study will inspire the development of more effective methods for the removal of the BCG, capable of isolating and attenuating artifact occurrences while preserving true neuronal activity.

---

**Supplementary Figure 1.**
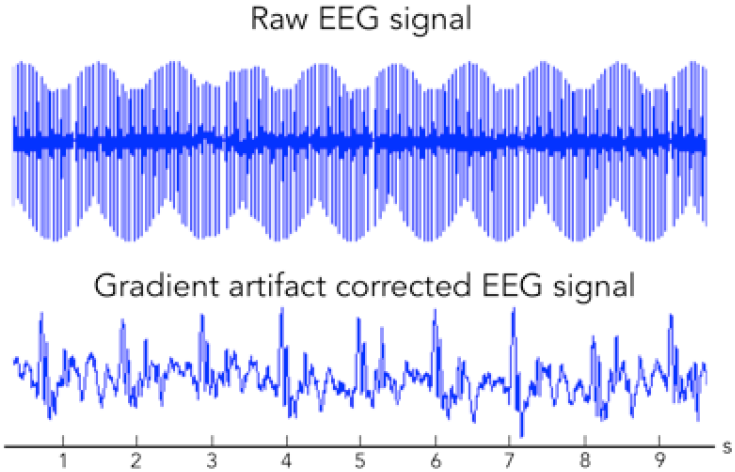
Representative EEG trace before and after the gradient artifact correction. The removal of the gradient artifact from the EEG signal permitted to identify BCG events.

**Supplementary Figure 2.**
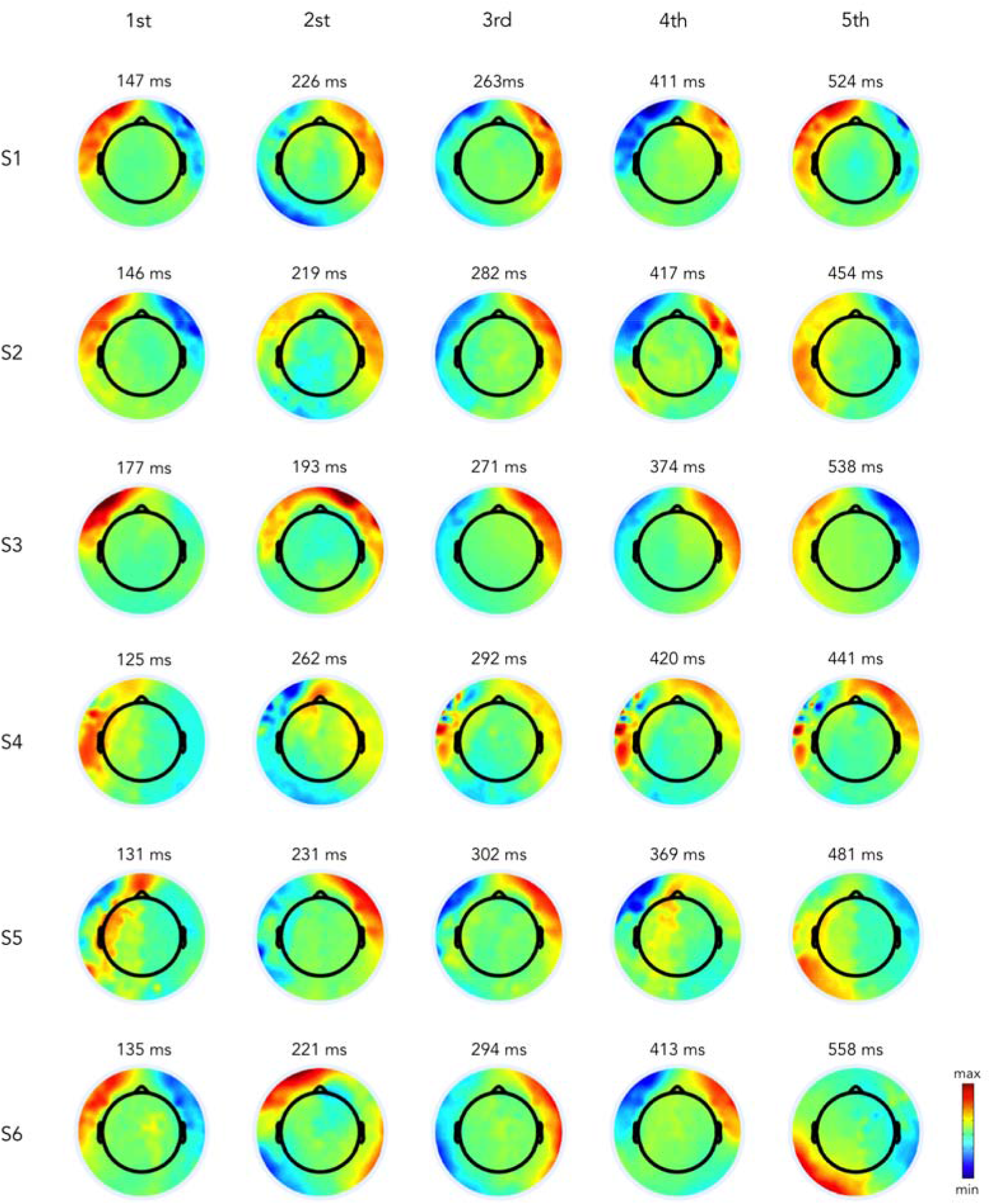
Subject-level spatial maps for each of the five main BCG peaks. Topographic maps were extracted for each subject in correspondence to five different major peaks identified from the RMS plot.

## BIBLIOGRAPHY

Allen, P. J., Josephs, O., & Turner, R. (2000). A method for removing imaging artifact from continuous EEG recorded during functional MRI. Neuroimage, 12(2), 230–239. doi: 10.1006/nimg.2000.0599

Allen, P. J., Polizzi, G., Krakow, K., Fish, D. R., & Lemieux, L. (1998). Identification of EEG events in the MR scanner: the problem of pulse artifact and a method for its subtraction. Neuroimage, 8(3), 229–239. doi: 10.1006/nimg.1998.0361

Bonmassar, G., Purdon, P. L., Jaaskelainen, I. P., Chiappa, K., Solo, V., Brown, E. N., & Belliveau, J. W. (2002). Motion and ballistocardiogram artifact removal for interleaved recording of EEG and EPs during MRI. Neuroimage, 16(4), 1127–1141.

Comon P. (1994). Independent component analysis - A new concept? Signal Processing. 36, 287–314.

Debener, S., Mullinger, K. J., Niazy, R. K., & Bowtell, R. W. (2008). Properties of the ballistocardiogram artefact as revealed by EEG recordings at 1.5, 3 and 7 T static magnetic field strength. Int J Psychophysiol, 67(1), 189–199. doi: 10.1016/j.ijpsych0.2007.05.015

Debener, S., Strobel, A., Sorger, B., Peters, J., Kranczioch, C., Engel, A. K., & Goebel, R. (2007). Improved quality of auditory event-related potentials recorded simultaneously with 3-T fMRI: removal of the ballistocardiogram artefact. Neuroimage, 34(2), 587–597. doi: 10.1016/j.neuroimage.2006.09.031

Debener, S., Ullsperger, M., Siegel, M., Fiehler, K., von Cramon, D. Y., & Engel, A. K. (2005). Trial-by-trial coupling of concurrent electroencephalogram and functional magnetic resonance imaging identifies the dynamics of performance monitoring. J Neurosci, 25(50), 11730–11737. doi: 10.1523/JNEUROSCI.3286-05.2005

Delorme, A., & Makeig, S. (2004). EEGLAB: an open source toolbox for analysis of single-trial EEG dynamics including independent component analysis. J Neurosci Methods, 134(1), 9–21. doi: 10.1016/j,jneumeth.2003.10.009

Grouiller F, Jorge J, Pittau F, van der Zwaag W, Iannotti GR, Michel CM, Vuilliemoz S, Vargas MI, Lazeyras F. (2016). Presurgical brain mapping in epilepsy using simultaneous EEG and functional MRI at ultra-high field: feasibility and first results. Magma; 29(3):605–16

Grouiller, F., Vercueil, L., Krainik, A., Segebarth, C, Kahane, P., & David, O. (2007). A comparative study of different artefact removal algorithms for EEG signals acquired during functional MRI. Neuroimage, 38(1), 124–137. doi: 10.1016/j.neuroimage.2007.07.025

Iannotti, G. R., Pittau, F., Michel, C. M., Vulliemoz, S., & Grouiller, F. (2015). Pulse artifact detection in simultaneous EEG-fMRI recording based on EEG map topography. Brain Topogr, 28(1), 21–32. doi: 10.1007/s10548-014-0409-z

Krishnaswamy P, Bonmassar G, Poulsen C, Pierce ET, Purdon PL, Brown EN. (2016). Reference-free removal of EEG-fMRI ballistocardiogram artifacts with harmonic regression. NeuroImage; 128:398–412

LeVan P, Maclaren J, Herbst M, Sostheim R, Zaitsev M, Hennig J. (2013). Ballistocardiographic artifact removal from simultaneous EEG-fMRI using an optical motion-tracking system. NeuroImage; 75:1–11

Liu Q, Balsters JH, Baechinger M, van der Groen O, Wenderoth N, Mantini D. (2015). Estimating a neutral reference for electroencephalographic recordings: the importance of using a high-density montage and a realistic head model. Journal of neural engineering; 12(5):056012

Mantini, D., Perrucci, M. G., Cugini, S., Ferretti, A., Romani, G. L., & Del Gratta, C. (2007). Complete artifact removal for EEG recorded during continuous fMRI using independent component analysis. Neuroimage, 34(2), 598–607. doi: 10.1016/j.neuroimage.2006.09.037

Mantini, D., Perrucci, M. G., Del Gratta, C, Romani, G. L., & Corbetta, M. (2007). Electrophysiological signatures of resting state networks in the human brain. Proc Natl Acad SciUSA, 104(32), 13170–13175. doi: 10.1073/pnas.0700668104

Mantini D, Marzetti L, Corbetta M, Romani GL, Del Gratta C. (2010). Multimodal integration of fMRI and EEG data for high spatial and temporal resolution analysis of brain networks. Brain topography; 23(2):150–8

Masterton, R. A., Abbott, D. F., Fleming, S. W., & Jackson, G. D. (2007). Measurement and reduction of motion and ballistocardiogram artefacts from simultaneous EEG and fMRI recordings. Neuroimage, 37(1), 202–211. doi: 10.1016/j.neuroimage.2007.02.060

McAvoy M, Mitra A, Tagliazucchi E, Laufs H, Raichle ME. (2017). Mapping visual dominance in human sleep. Neuroimage; 150:250–61

Mullinger, K. J., Havenhand, J., & Bowtell, R. (2013). Identifying the sources of the pulse artefact in EEG recordings made inside an MR scanner. Neuroimage, 71, 75–83. doi: 10.1016/j.neuroimage.2012.12.070

Müri, R. M., Felblinger, J., Rosler, K. M., Jung, B., Hess, C. W., & Boesch, C. (1998). Recording of electrical brain activity in a magnetic resonance environment: distorting effects of the static magnetic field. Magn Reson Med, 39(1), 18–22

Neuner I, Arrubla J, Felder J, Shah NJ. (2014). Simultaneous EEG-fMRI acquisition at low, high and ultra-high magnetic fields up to 9.4 T: perspectives and challenges. Neuroimage; 102 Pt 1: 71–9

Niazy, R. K., Beckmann, C. F., lannetti, G. D., Brady, J. M., & Smith, S. M. (2005). Removal of FMRI environment artifacts from EEG data using optimal basis sets. Neuroimage, 28(3), 720–737. doi: 10.1016/j.neuroimage.2005.06.067

Oh, S. S., Han, Y., Lee, J., Yun, S. D., Kang, J. K., Lee, E. M., Yoon, H.W., Chung, J.Y., Park, H. (2014). A pulse artifact removal method considering artifact variations in the simultaneous recording of EEG and fMRI. Neurosci Res, 81-82, 42–50. doi: 10.1016/j.neures.2014.01.008

Srivastava, G., Crottaz-Herbette, S., Lau, K. M., Glover, G. H., & Menon, V. (2005). ICA-based procedures for removing ballistocardiogram artifacts from EEG data acquired in the MRI scanner. Neuroimage, 24(1), 50–60. doi: 10.1016/j.neuroimage.2004.09.041

Yan, W. X., Mullinger, K. J., Geirsdottir, G. B., & Bowtell, R. (2010). Physical modeling of pulse artefact sources in simultaneous EEG/fMRI. Hum Brain Mapp, 31(4), 604–620. doi: 10.1002/hbm.20891

